## Supplementary Information for "C9orf72 polyPR directly binds to various nuclear transport components"

### 1 Coarse-grained 1-BPA force field

#### 1.1 PolyPR–polyPR interaction

We use the 1-bead-per-amino-acid (1BPA) force field [1, 2] for polyPR-polyPR interactions. The 1BPA force field has been previously used to study intrinsically disordered FG-Nups and dipeptide repeat proteins (DPRs) [3, 4]. The bonded interactions, i.e. the bending and torsion potentials, in this force field are residue and sequence specific [1]. The attractive hydrophobic and repulsive hydrophilic interactions between different residues in this force field are represented by:

$$\phi_{\mathrm{hp}} = \begin{cases} \varepsilon_{\mathrm{rep}} \left(\frac{\sigma}{r}\right)^8 - \varepsilon_{ij} \left[\frac{4}{3}\left(\frac{\sigma}{r}\right)^6 - \frac{1}{3}\right] & r \leq \sigma \\ (\varepsilon_{\mathrm{rep}} - \varepsilon_{ij}) \left(\frac{\sigma}{r}\right)^8 & r \geq \sigma, \end{cases}$$

where $\varepsilon_{ij} = \varepsilon_{\mathrm{hp}} \sqrt{(\varepsilon_i \varepsilon_j)^{0.27}}$ is the strength of the interaction for each pair of amino acids ($i$,$j$), $r$ is the distance between beads $i$ and $j$, and $\sigma = 0.6$ nm. The values of $\varepsilon_{\mathrm{hp}}$ and $\varepsilon_{\mathrm{rep}}$ are 13 and 10 kJ/mol, respectively. The relative hydrophobic strength values ($\varepsilon_i \in [0,1]$) of the different amino acids are listed in Table S1 [2]. The hydrophobic strength values of charged residues are slightly increased in line with our recent work [4].

The electrostatic interactions between charged residues are described by the modified Coulomb law:

$$\phi_{\mathrm{elec}} = \frac{q_i q_j}{4\pi\varepsilon_0 \varepsilon_r(r) r} e^{-\kappa r},$$

where $\varepsilon_r(r) = S_{\mathrm{s}} \left[1 - \frac{r^2}{z^2} \frac{e^{r/z}}{(e^{r/z}-1)^2}\right]$ is the distance-dependent dielectric constant of the solvent with $S_s = 80$ and $z = 0.25$ nm. The value of the Debye screening coefficient, $\kappa$, is 1 nm$^{-1}$ for monovalent salt concentration $C_{\mathrm{salt}}$ = 100 mM, and 1.5 nm$^{-1}$ for $C_{\mathrm{salt}}$ = 200 mM.

For the interactions between the residues within the disordered regions of transport components we also use the 1BPA force field featuring $\phi_{\mathrm{hp}}$ and $\phi_{\mathrm{elec}}$ as described above.

#### 1.2 PolyPR interaction with transport components

Poly-PR has been shown to bind to several importins in *in vitro* experiments [5]. However, no binding has been observed for the more hydrophobic DPRs, i.e. poly-GA and poly-GP [5]. These observations highlight the importance of Arginine in driving the binding between poly-PR and NCT components. At physiological salt concentrations, Arginine mainly engages in electrostatic and cation-pi interactions. For the polyPR interaction with transport components, we use the same electrostatic potential ($\phi_{\mathrm{elec}}$) as described in the previous section. To take into account the cation-pi interactions between Arginine (in polyPR) and the aromatic residues Phenylalanine, Tyrosine and Tryptophan (in the transport components), we use an 8-6 Lennard-Jones (LJ) potential that replaces $\phi_{\mathrm{hp}}$ for the RF, RY, and RW interactions:

$$\phi_{\mathrm{cp},ij}(r) = \varepsilon_{\mathrm{cp},ij}\left[3\left(\frac{r_{\mathrm{m}}}{r}\right)^{8} - 4\left(\frac{r_{\mathrm{m}}}{r}\right)^{6}\right],$$

where $r_{\mathrm{m}} = 0.45$ nm is the distance at which the $\phi_{\mathrm{cp},ij}$ reaches its minimum value, and $\varepsilon_{\mathrm{cp},ij}$ is a pair-dependent cation-pi energy taken from [6] for different combinations of cation-pi interactions, as listed below

| Cation-pi pair | $\boldsymbol{\varepsilon}_{\mathbf{cp,RF}}$ | $\boldsymbol{\varepsilon}_{\mathbf{cp,RY}}$ | $\boldsymbol{\varepsilon}_{\mathbf{cp,RW}}$ | $\boldsymbol{\varepsilon}_{\mathbf{cp,KF}}$ | $\boldsymbol{\varepsilon}_{\mathbf{cp,KY}}$ | $\boldsymbol{\varepsilon}_{\mathbf{cp,KW}}$ |
|---|---|---|---|---|---|---|
| Energy (kJ/mol) | 4.30 | 5.00 | 6.70 | 1.79 | 3.13 | 4.26 |

For the hydrophilic/hydrophobic interactions between polyPR and the rest of the transport component residues, including the disordered regions, we use $\phi_{\mathrm{hp}}$ with $\varepsilon_{ij} = 10$ kJ/mol which leads to an excluded volume potential that vanishes at $r = 0.6$ nm.

#### 2 Supplementary figures

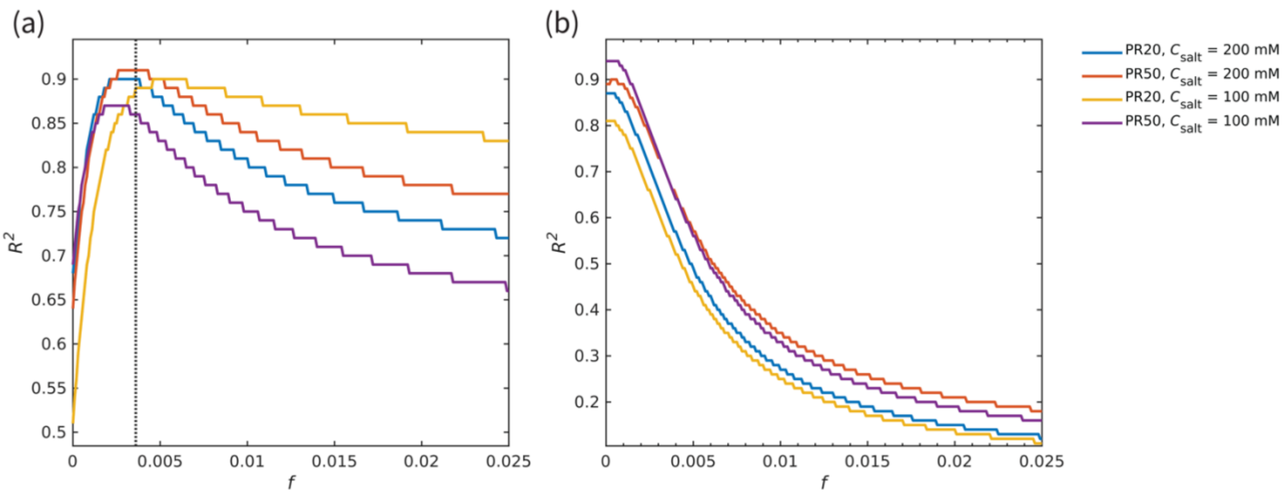

**Figure S1**: The quality of fit ($R^2$) for different values of the dimensionless parameter $f$ in Eq. (1) for (a) the transport components shown in figure 1, excluding the specific exporters of Impα, CAS and Cse1, and (b) the Kapβ set (studied here [6]) together with CAS and Cse1. The results are shown for PR20, 50 and the two salt concentrations used in this study. In (a) the vertical dotted line in the left panel shows the value of $f = 0.0036$ for the best fit based on the four curves.

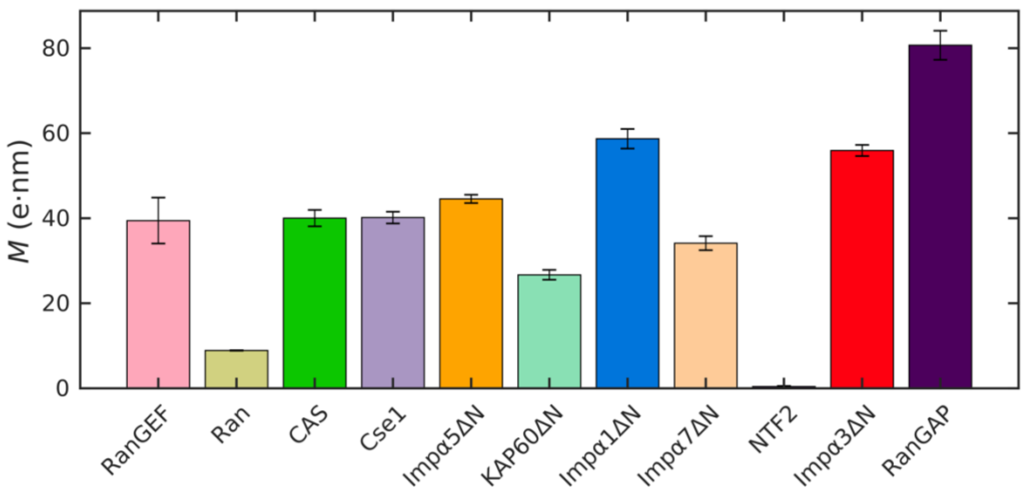

**Figure S2**: Dipole moment of the different transport components shown in figure 1.

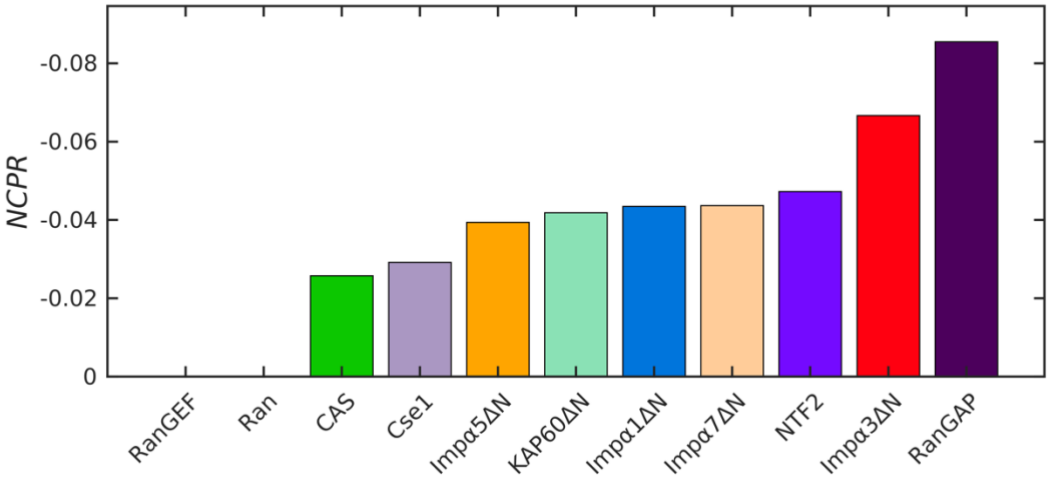

**Figure S3**: Net charge per residue of the different transport components shown in figure 1.

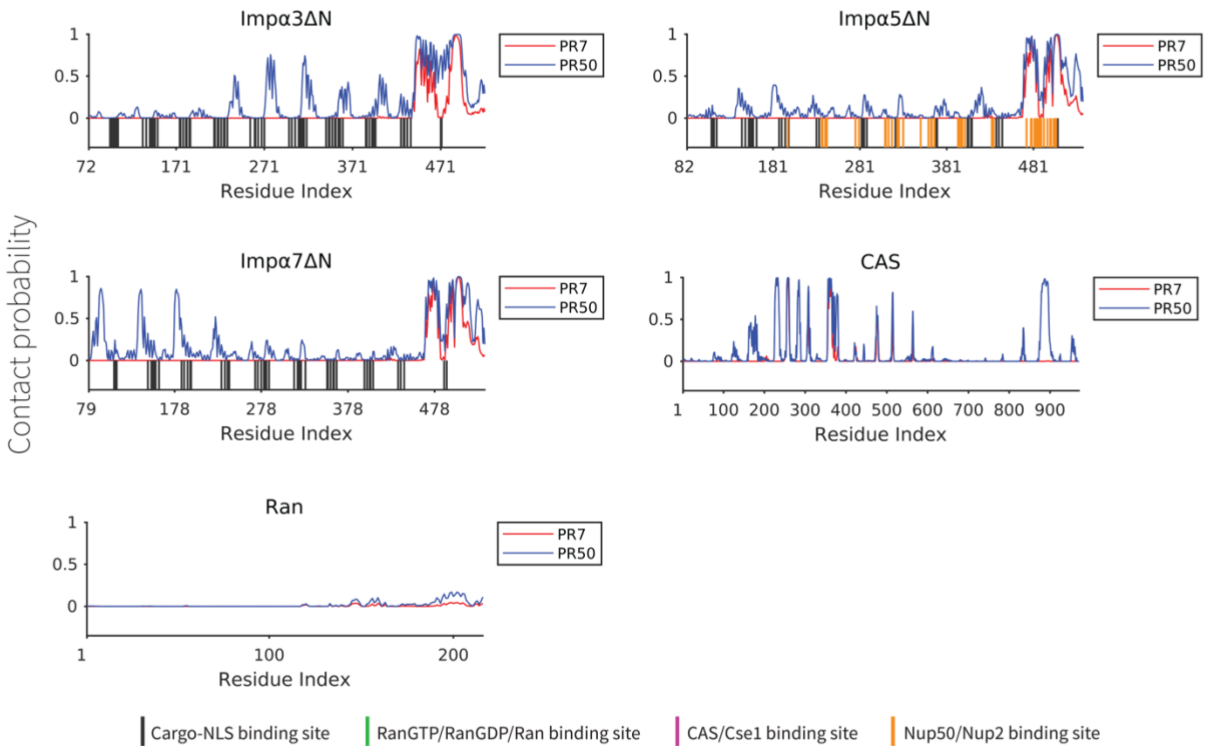

**Figure S4**: The contact probability for each residue in the sequence of transport components interacting with polyPR. The plot displays the contact probability for six transport components: Impα3ΔN, Impα5ΔN, Impα7ΔN, CAS, and Ran at a salt concentration of 100 mM.

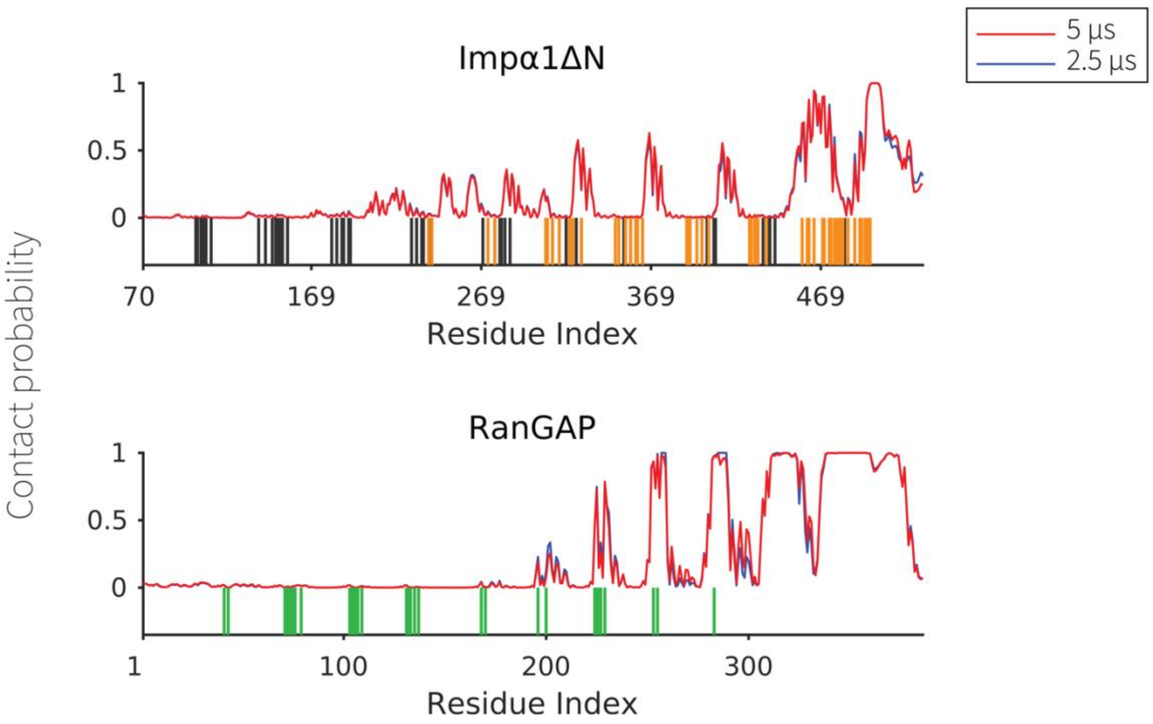

**Figure S5:** The contact probability for each residue in the sequence of transport components interacting with PR50. The plot displays the contact probability calculated from simulations performed for 2.5 and 5 μs for Impα1ΔN and RanGAP, showing convergence of the computations.

### 3 Supplementary tables

**Table S1** Relative hydrophobic strength values of the different amino acids [2, 4].

| Amino acid | $\varepsilon_i$ | Amino acid | $\varepsilon_i$ |
|---|---|---|---|
| A | 0.7 | L | 1 |
| R | 0.005 | K | 0.005 |
| N | 0.33 | M | 0.78 |
| D | 0.005 | F | 1 |
| C | 0.68 | P | 0.65 |
| Q | 0.64 | S | 0.45 |
| E | 0.005 | T | 0.51 |
| G | 0.41 | W | 0.96 |
| H | 0.53 | Y | 0.82 |
| I | 0.98 | V | 0.94 |

**Table S2** Information about the transport components used in this study.

| Transport component | Uniprot ID | Organism | PDB | Seq. length | Res. in the model | Binding partners of each transport component (PDB code) |
|---|---|---|---|---|---|---|
| **Impα1 (KPNA2)** | P52292 | Human | 4e4v | 529 | 70-529 | Protein cargoes: yeast CBP80 cNLS (3uky), Dengue 3 NS5 C-terminal NLS peptide (5fc8), Dengue 2 NS5 C-terminal NLS peptide (5hhg), Zika MR766 NLS (5w41), Nipah virus W protein C-terminus (6bw0), Hendra virus W protein C-terminus (6bw1), XRCC1 NLS peptide (5e6q), Influenza PB2 NLS (4uaf), PARP-2 NLS (5d5k), TPX2 (3knd), Pom121 NLS (4yi0), SART3/TIP110 NLS (5ctt), MAL RPEL (3tpm) Nup50: Nup50 (2c1m) |
| **Impα3 (KPNA4)** | O00629 | Human | 6bvz | 521 | 72-521 | Protein cargoes: Hendra virus W protein C-terminus crystal forms 1,2,3 (6bw9, 6bwa, 6bwb), Nipah virus W protein C-terminus (6bvv), RCC1(5tbk), Influenza A PB2 NLS (4uae), N-terminal fragment of RanBP3 (5xzx) |
| **Impα5 (KPNA1)** | P52294 | Human | AlphaFold | 538 | 82-538 | Protein cargoes: C-terminal domain of Influenza virus PB2 subunit (2jdq) Nup50: N-terminus of Nup50 (3tj3) |
| **Impα7 (KPNA6)** | O60684 | Human | AlphaFold | 536 | 79-536 | Protein cargoes: Influenza A PB2 NLS (4uad) |
| **KAP60** | Q02821 | *S. cerevisiae* | 1bk5 | 541 | 89-541 | Protein cargoes: SUMO Protease Ulp1p (5h2w), INM protein Heh1 (4xzr), INM protein Heh2 (4pvz), yeast RCC1 (5t94) Nup2 (yeast Nup50): Nup2p N-terminal fragment (1un0), Nup2 (2c1t) Cse1: CSE1 (1wa5) RanGTP: RanGTP (1wa5) |
| **CAS** | P55060 | Human | AlphaFold | 971 | 1-971 | N.A. |
| **Cse1** | P33307 | *S. cerevisiae* | 1z3h | 960 | 1-960 | Impα: Cse1 (1wa5) RanGTP: RanGTP (1wa5) |
| **RanGEF (RCC1)** | P18754 | Human | 1a12 | 421 | 1-421 | Ran protein: Ran (1i2m) |
| **RanGAP** | P41391 | *S. pombe* | AlphaFold | 386 | 1-386 | RanGTP: RanGPPNHP (1k5d) |
| **NTF2*** | P61972 P61970 | Rat Human | 1oun | 127 | 1-127 | RanGDP: RanGDP (5bxq, 1a2k) |
| **Ran** | P62826 | Human | 2mmc | 216 | 1-216 | N.A. |

* The amino acid sequence of NTF2 Rat is identical to the human version.

*NTF2 is a homodimer and the CG model contains 254 residues in total.

**Table S3** Sequences of transport components used for the CG modeling. See also column six of Table S2 for more information about the transport components used in this study.

| NTR name | Amino acid sequence |
|---|---|
| Impα1ΔN | NQGTVNWSVDDIVKGINSSNVENQLQATQAARKLLSREKQPPIDNIIRA<br>LIPKFVSFLGRTDCSPIQFESAWALTNIASGTSEQTKAVVDGGAIPAFI<br>LLASPHAHISEQAVWALGNIAGDGSVFRDLVIKYGAVDPLLALLAVPDM<br>SLACGYLRNLTWTLSNLCRNKNPAPPIDAVEQILPTLVRLLHHDDPEVL<br>DTCWAISYLTDGPNERIGMVVKTGVVPQLVKLLGASELPIVTPALRAIG<br>IVTGTDEQTQVVIDAGALAVFPSLLTNPKTNIQKEATWTMSNITAGRQD<br>IQQVVNHGLVPFLVSVLSKADFKTQKEAVWAVTNYTSGGTVEQIVYLVH<br>GIIEPLMNLLTAKDTKIILVILDAISNIFQAAEKLGETEKLSIMIEECG<br>LDKIEALQNHENESVYKASLSLIEKYFSVEEEEDQNVVPETTSEGYTFQ<br>QDGAPGTFNF |
| Impα3ΔN | SLEAIVQNASSDNQGIQLSAVQAARKLLSSDRNPPIDDLIKSGILPILV<br>CLERDDNPSLQFEAAWALTNIASGTSEQTQAVVQSNAVPLFLRLLHSPH<br>NVCEQAVWALGNIIGDGPQCRDYVISLGVVKPLLSFISPSIPITFLRNV<br>WVMVNLCRHKDPPPPMETIQEILPALCVLIHHTDVNILVDTVWALSYLT<br>AGNEQIQMVIDSGIVPHLVPLLSHQEVKVQTAALRAVGNIVTGTDEQTQ<br>VLNCDALSHFPALLTHPKEKINKEAVWFLSNITAGNQQQVQAVIDANLV<br>MIIHLLDKGDFGTQKEAAWAISNLTISGRKDQVAYLIQQNVIPPFCNLL<br>VKDAQVVQVVLDGLSNILKMAEDEAETIGNLIEECGGLEKIEQLQNHEN<br>DIYKLAYEIIDQFFSSDDIDEDPSLVPEAIQGGTFGFNSSANVPTEGFQ |
| Impα5ΔN | VITSDMIEMIFSKSPEQQLSATQKFRKLLSKEPNPPIDEVISTPGVVAR<br>VEFLKRKENCTLQFESAWVLTNIASGNSLQTRIVIQAGAVPIFIELLSS<br>FEDVQEQAVWALGNIAGDSTMCRDYVLDCNILPPLLQLFSKQNRLTMTR<br>AVWALSNLCRGKSPPPEFAKVSPCLNVLSWLLFVSDTDVLADACWALSY<br>SDGPNDKIQAVIDAGVCRRLVELLMHNDYKVVSPALRAVGNIVTGDDIQ<br>QVILNCSALQSLLHLLSSPKESIKKEACWTISNITAGNRAQIQTVIDAN<br>FPALISILQTAEFRTRKEAAWAITNATSGGSAEQIKYLVELGCIKPLCD<br>LTVMDSKIVQVALNGLENILRLGEQEAKRNGTGINPYCALIEEAYGLDK<br>EFLQSHENQEIYQKAFDLIEHYFGTEDEDSSIAPQVDLNQQQYIFQQCE<br>PMEGFQL |
| Impα7ΔN | SVITREMVEMLFSDDSDLQLATTQKFRKLLSKEPSPPIDEVINTPRVVD<br>FVEFLKRNENCTLQFEAAWALTNIASGTSQQTKIVIEAGAVPIFIELLN<br>DFEDVQEQAVWALGNIAGDSSVCRDYVLNCSILNPLLTLLTKSTRLTMT<br>NAVWALSNLCRGKNPPPEFAKVSPCLPVLSRLLFSSDSDLLADACWALS<br>LSDGPNEKIQAVIDSGVCRRLVELLMHNDYKVASPALRAVGNIVTGDDI<br>TQVILNCSALPCLLHLLSSPKESIRKEACWTISNITAGNRAQIQAVIDA<br>IFPVLIEILQKAEFRTRKEAAWAITNATSGGTPEQIRYLVSLGCIKPLC<br>LLTVMDSKIVQVALNGLENILRLGEQEGKRSGSGVNPYCGLIEEAYGLD<br>IEFLQSHENQEIYQKAFDLIEHYFGVEDDDSSLAPQVDETQQQFIFQQP<br>APMEGFQL |

| | |
|---|---|
| KAP60ΔN | LPQMTQQLNSDDMQEQLSATVKFRQILSREHRPPIDVVIQAGVVPRLVE<br>MRENQPEMLQLEAAWALTNIASGTSAQTKVVVDADAVPLFIQLLYTGSV<br>VKEQAIWALGNVAGDSTDYRDYVLQCNAMEPILGLFNSNKPSLIRTATW<br>LSNLCRGKKPQPDWSVVSQALPTLAKLIYSMDTETLVDACWAISYLSDG<br>QEAIQAVIDVRIPKRLVELLSHESTLVQTPALRAVGNIVTGNDLQTQVV<br>NAGVLPALRLLLSSPKENIKKEACWTISNITAGNTEQIQAVIDANLIPP<br>VKLLEVAEYKTKKEACWAISNASSGGLQRPDIIRYLVSQGCIKPLCDLL<br>IADNRIIEVTLDALENILKMGEADKEARGLNINENADFIEKAGGMEKIF<br>CQQNENDKIYEKAYKIIETYFGEEEDAVDETMAPQNAGNTFGFGSNVNQ<br>FNFN |
| CAS | MELSDANLQTLTEYLKKTLDPDPAIRRPAEKFLESVEGNQNYPLLLLTL<br>EKSQDNVIKVCASVTFKNYIKRNWRIVEDEPNKICEADRVAIKANIVHL<br>LSSPEQIQKQLSDAISIIGREDFPQKWPDLLTEMVNRFQSGDFHVINGV<br>RTAHSLFKRYRHEFKSNELWTEIKLVLDAFALPLTNLFKATIELCSTHA<br>DASALRILFSSLILISKLFYSLNFQDLPEFFEDNMETWMNNFHTLLTLD<br>KLLQTDDEEEAGLLELLKSQICDNAALYAQKYDEEFQRYLPRFVTAIWN<br>LVTTGQEVKYDLLVSNAIQFLASVCERPHYKNLFEDQNTLTSICEKVIV<br>NMEFRAADEEAFEDNSEEYIRRDLEGSDIDTRRRAACDLVRGLCKFFEG<br>VTGIFSGYVNSMLQEYAKNPSVNWKHKDAAIYLVTSLASKAQTQKHGIT<br>ANELVNLTEFFVNHILPDLKSANVNEFPVLKADGIKYIMIFRNQVPKEH<br>LVSIPLLINHLQAESIVVHTYAAHALERLFTMRGPNNATLFTAAEIAPF<br>EILLTNLFKALTLPGSSENEYIMKAIMRSFSLLQEAIIPYIPTLITQLT<br>KLLAVSKNPSKPHFNHYMFEAICLSIRITCKANPAAVVNFEEALFLVFT<br>ILQNDVQEFIPYVFQVMSLLLETHKNDIPSSYMALFPHLLQPVLWERTG<br>IPALVRLLQAFLERGSNTIASAAADKIPGLLGVFQKLIASKANDHQGFY<br>LNSIIEHMPPESVDQYRKQIFILLFQRLQNSKTTKFIKSFLVFINLYCI<br>YGALALQEIFDGIQPKMFGMVLEKIIIPEIQKVSGNVEKKICAVGITKL<br>TECPPMMDTEYTKLWTPLLQSLIGLFELPEDDTIPDEEHFIDIEDTPGY<br>TAFSQLAFAGKKEHDPVGQMVNNPKIHLAQSLHKLSTACPGRVPSMVST<br>LNAEALQYLQGYLQAASVTLL |
| Cse1 | MSDLETVAKFLAESVIASTAKTSERNLRQLETQDGFGLTLLHVIASTNL<br>LSTRLAGALFFKNFIKRKWVDENGNHLLPANNVELIKKEIVPLMISLPN<br>LQVQIGEAISSIADSDFPDRWPTLLSDLASRLSNDDMVTNKGVLTVAHS<br>FKRWRPLFRSDELFLEIKLVLDVFTAPFLNLLKTVDEQITANENNKASL<br>ILFDVLLVLIKLYYDFNCQDIPEFFEDNIQVGMGIFHKYLSYSNPLLED<br>DETEHASVLIKVKSSIQELVQLYTTRYEDVFGPMINEFIQITWNLLTSI<br>NQPKYDILVSKSLSFLTAVTRIPKYFEIFNNESAMNNITEQIILPNVTL<br>EEDVELFEDDPIEYIRRDLEGSDTDTRRRACTDFLKELKEKNEVLVTNI<br>LAHMKGFVDQYMSDPSKNWKFKDLYIYLFTALAINGNITNAGVSSTNNL<br>NVVDFFTKEIAPDLTSNNIPHIILRVDAIKYIYTFRNQLTKAQLIELMP<br>LATFLQTDEYVVVYTYAAITIEKILTIRESNTSPAFIFHKEDISNSTEIL<br>KNLIALILKHGSSPEKLAENEFLMRSIFRVLQTSEDSIQPLFPQLLAQF<br>EIVTIMAKNPSNPRFTHYTFESIGAILNYTQRQNLPLLVDSMMPTFLTV<br>SEDIQEFIPYVFQIIAFVVEQSATIPESIKPLAQPLLAPNVWELKGNIP<br>VTRLLKSFIKTDSSIFPDLVPVLGIFQRLIASKAYEVHGFDLLEHIMLL<br>DMNRLRPYIKQIAVLLLQRLQNSKTERYVKKLTVFFGLISNKLGSDFLI<br>FIDEVQDGLFQQIWGNFIITTLPTIGNLLDRKIALIGVLNMVINGQFFQ<br>KYPTLISSTMNSIIETASSQSIANLKNDYVDLDNLEEISTFGSHFSKLV |

| | |
|---|---|
| | ISEKPFDPLPEIDVNNGVRLYVAEALNKYNAISGNTFLNTILPQLTQEN<br>VKLNQLLVGN |
| RanGEF<br>(RCC1) | MSPKRIAKRRSPPADAIPKSKKVKVSHRSHSTEPGLVLTLGQGDVGQLG<br>GENVMERKKPALVSIPEDVVQAEAGGMHTVCLSKSGQVYSFGCNDEGAL<br>RDTSVEGSEMVPGKVELQEKVVQVSAGDSHTAALTDDGRVFLWGSFRDN<br>GVIGLLEPMKKSMVPVQVQLDVPVVKVASGNDHLVMLTADGDLYTLGCG<br>QGQLGRVPELFANRGGRQGLERLLVPKCVMLKSRGSRGHVRFQDAFCGA<br>FTFAISHEGHVYGFGLSNYHQLGTPGTESCFIPQNLTSFKNSTKSWVGF<br>GGQHHTVCMDSEGKAYSLGRAEYGRLGLGEGAEEKSIPTLISRLPAVSS<br>ACGASVGYAVTKDGRVFAWGMGTNYQLGTGQDEDAWSPVEMMGKQLENR<br>VLSVSSGGQHTVLLVKDKEQS |
| RanGAP | MSRFSIEGKSLKLDAITTEDEKSVFAVLLEDDSVKEIVLSGNTIGTEAA<br>WLSENIASKKDLEIAEFSDIFTGRVKDEIPEALRLLLQALLKCPKLHTV<br>LSDNAFGPTAQEPLIDFLSKHTPLEHLYLHNNGLGPQAGAKIARALQEL<br>VNKKAKNAPPLRSIICGRNRLENGSMKEWAKTFQSHRLLHTVKMVQNGI<br>PEGIEHLLLEGLAYCQELKVLDLQDNTFTHLGSSALAIALKSWPNLREL<br>LNDCLLSARGAAAVVDAFSKLENIGLQTLRLQYNEIELDAVRTLKTVID<br>KMPDLLFLELNGNRFSEEDDVVDEIREVFSTRGRGELDELDDMEELTDE<br>EEDEEEEAESQSPEPETSEEEKEDKELADELSKAHI |
| NTF2 | MGDKPIWEQIGSSFIQHYYQLFDNDRTQLGAIYIDASCLTWEGQQFQGK<br>AIVEKLSSLPFQKIQHSITAQDHQPTPDSCIISMVVGQLKADEDPIMGF<br>QMFLLKNINDAWVCTNDMFRLALHNFG |
| Ran | MAAQGEPQVQFKLVLVGDGGTGKTTFVKRHLTGEFEKKYVATLGVEVHP<br>VFHTNRGPIKFNVWDTAGQEKFGGLRDGYYIQAQCAIIMFDVTSRVTYK<br>VPNWHRDLVRVCENIPIVLCGNKVDIKDRKVKAKSIVFHRKKNLQYYDI<br>AKSNYNFEKPFLWLARKLIGDPNLEFVAMPALAPPEVVMDPALAAQYEH<br>LEVAQTTALPDEDDDL |

**Table S4:** Parameters for the best linear fits shown in figure 2.

| | figure 2a ($\boldsymbol{ax+b}$) | | | | figure 2b ($\boldsymbol{ax+b}$) | | | |
|---|---|---|---|---|---|---|---|---|
| | PR20 | | PR50 | | PR20 | | PR50 | |
| | $a$ | $b$ | $a$ | $b$ | $a$ | $b$ | $a$ | $b$ |
| $\boldsymbol{C_{\mathrm{salt}} = 200}$ mM | -0.0856 | -0.0027 | -0.0481 | -0.0016 | -0.1216 | -0.0014 | -0.0866 | -0.0008 |
| $\boldsymbol{C_{\mathrm{salt}} = 100}$ mM | -0.0955 | -0.0001 | -0.0638 | -0.0002 | -0.0786 | 0.0027 | -0.0754 | 0.0015 |
